# Combinatorial approach for complex disorder prediction: Case study of neurodevelopmental disorders

**DOI:** 10.1101/129775

**Authors:** Linh Huynh, Fereydoun Hormozdiari

## Abstract

Early prediction of complex disorders (e.g., autism and other neurodevelopmental disorders) is one of the fundamental goals of precision medicine and personalized genomics. An early prediction of complex disorders can have a significant impact on increasing the effectiveness of interventions and treatments in improving the prognosis and, in many cases, enhancing the quality of life in the affected patients. Considering the genetic heritability of neurodevelopmental disorders, we are proposing a novel framework for utilizing rare coding variation for early prediction of these disorders in subset of affected samples. We provide a novel formulation for the **U**ltra-**A**ccurate **D**isorder **P**rediction (UADP) problem and develop a combinatorial framework for solving this problem. The primary goal of this framework, denoted as Odin (**O**racle for **DI**sorder predictio**N**), is to make prediction for a subset of affected cases while having very low false positive rate prediction for unaffected samples. Note that in the Odin framework we will take advantage of the available functional information (e.g., pairwise coexpression of genes during brain development) to increase the prediction power beyond genes with recurrent variants. Application of our method accurately recovers an additional 8% of autism cases without a sever variant in a known recurrent mutated genes with a less than 1% false positive rate. Furthermore, Odin predicted a set of 391 genes that severe variants in these genes can cause autism or other developmental delay disorders. Odin is publicly available at https://github.com/HormozdiariLab/Odin ^†^

## 1 Introduction

The start of the genomics era and sequencing of the first human genome over a decade ago promised significant benefits to public health [1]. These include the potential capability of early detection, pinpointing the causes, and developing novel treatments and therapeutics for most diseases. The sequencing of the human genome has dramatically accelerated biomedical research; however, even a decade after publication, progress has been slow in truly unlocking the promise of genetics and genomics in direct application to human health and disease. Notably, the translation of genetic discoveries into actionable items in medicine has not achieved the promised potential. One of the main challenges lies in the fact that discovering the exhaustive set of causative variants for most diseases, except some monogenic Mendelian disorders, has proven to be an elusive and unmet objective [2–4].

One of the first questions that we need to answer regarding any disease of interest is to calculate the contribution of genetics to the etiology of the disease. The primary metric used for calculating the contribution of genetics to any trait, including diseases, is denoted as genetic heritability (0 ≤ *h*^2^ ≤ 1) [5, 6]. However, in many complex diseases, the calculated genetic heritability is by far higher than the fraction of cases that can be explained or predicted by the observed genetic variants. This gap is known as the “missing heritability” problem and is one of the main hindrances in not only building early prediction models for complex disorders but also developing novel treatments [7, 8].

Autism spectrum disorder (ASD) is an umbrella term used to describe a set of neurodevelopmental disorders having a wide range of symptoms from lack of social interaction, difficulty in communication/language, repetitive behavior, and in many cases intellectual disability (ID) (i.e., having an IQ < 70) [9]. ASD is typically diagnosed around the age of two and is estimated to affect over 1 in 68 children (1.5% of all children). There is a well-known sex bias in ASD as there are four times more male children affected with ASD than female children. Twin study comparisons have shown that genetics play a major role in ASD, and researchers have estimated the heritability of ASD to be one of the highest among complex diseases (0.5 ≤ *h*^2^ ≤ 0.8) [10, 11].

It is becoming apparent that early treatment and intervention can significantly improve the IQ, language skills, and social interactions in children affected with ASD [12–14]. Early diagnosis of ASD in young infants is challenging mainly due to the fact that most symptoms are not reliably detectable at a very young age and children tend to manifest a heterogeneous set of phenotypes with a diverse range of severity [15]. However, it is theoretically possible to make an accurate diagnosis of ASD or other neurodevelopmental disorders in **subset of children** before any symptoms appear (or even before the child is born) using (perinatal) genetic testing and genome sequencing [16]. Thus, building methods for *early prediction of ASD using genetic variation and other biomarkers* is extremely important and will have significant direct positive effects on public health.

The recent advances in high-throughput sequencing (HTS) technologies have given us the capability to sequence the whole genomes or exomes of many samples. For instance, the consortia focused on complex disorders, such as autism, ID, schizophrenia, and diabetes sequenced tens of thousands of cases [17–24]. Sequencing of samples with these complex disorders had produced genetic maps with tens of thousands of variants with most of them being rare (i.e., minor allele frequency - MAF < 0.05). Unfortunately, in most cases the effect of the variants found is not known (i.e., a variant of unknown significance) and for rare variants it is extremely hard to assign significance solely considering their frequency. In extreme cases of *de novo* variants (i.e., novel variants not inherited from the parents), the same exact variant will likely never be seen in any other sample. Thus, building models for early prediction of complex disorders need to be more sophisticated than just considering the frequency of variants in cases and controls [25].

There are some known syndromic subtypes of ASD with known genetic causes, such as Fragile X or Rett syndromes, which are the result of single-gene mutations (*FMR1* or *MECP2*, respectively) [26]. Furthermore, there are known rare, large recurrent copy number variations, such as the 16p11.2 deletion or Prader-Willi syndrome, which are known to cause ASD [27, 28]. Note that, these specific tests will predict ASD/ID for a small subset of affected cases with low false positive rate. However, in most cases of ASD, the exact cause of the disorder is not known and no accurate method or model for early prediction of ASD using genetic variants exists. Recently, several autism and intellectual disability sequencing consortia [17, 18] performed whole-exome sequencing on thousands of autism families (affected proband, unaffected sibling and parents) with the hope of finding causative variants in these samples. The enrichment of *de novo* variants in affected probands versus unaffected siblings has indicated that a significant fraction of ASD is the result of *de novo* and rare (MAF < 0.05) variants [17, 29, 30]. However, in many cases, it is not clear which *de novo* or rare variants are the real culprit(s) of the phenotype. As we do not expect to see the same *de novo* variant to appear in two different samples, it has proven useful to summarize the observed variants on the genes being affected. This simple approach has provided researchers with enough evidence to predict tens of novel ASD genes with high penetrance of **likely gene disruptive (LGD)** and missense variants [17]. However, these statistically significant genes only cover a fraction of ASD cases estimated to be caused by *de novo* or rare variants. Based on the twin studies, it is estimated that ASD and ID have a genetic heritability of over 0.5, while we can optimistically explain less than 0.2 fraction of the affected children [17, 29] based on observed common or rare genetic variants (including structural variation and copy number variation [31–33]).

It is estimated that hundreds of genes are involved in neurodevelopment and disruption in them can cause ASD or ID [29, 34–36]. The primary justification for such a high number of genes (high genetic heterogeneity) contributing to a similar disorder (i.e., ASD) is that most of these genes are members of only a few functional “modules” or pathways [37–43]. Thus, disruption of any of these functional modules results in a similar disorder mainly through interruption of normal neurodevelopment. It is being hypothesized that by using the functional relationship between these genes it is possible to find the causative variants.

### Complex disorder prediction using (rare) coding variants

As mentioned above, early prediction of complex disorders using genetic variation is one of the fundamental goals of personalized medicine. Currently, thousands of cases with neurodevelopmental disorders (e.g., ASD) have been studied using WES or targeted sequencing. Thus, we have a very rich set of rare and common coding variants found in samples with ASD and other neurodevelopmental disorders, which can be used for building early prediction models and methods. However, it is also important to realize the intrinsic limitations of rare coding variants in predicting ASD or other complex disorders. Notably, (i) most complex disorders have genetic heritability of significantly less than 1 (e.g., 0.5 < *h*^2^ < 0.8 for autism), (ii) noncoding variants, which significantly contribute to these disorders, are not found using WES, and (iii) (coding) variants alone do not have the power to *rule out* the possibility of being diagnosed of a complex disorder (such as autism) with very high accuracy. **Therefore, achieving accurate prediction for all (or even most) affected cases using solely the coding variants is theoretically not achievable. On the flip side, this also means, that we cannot confidently predict a sample as an unaffected control solely based on the observed (coding) variants, as other factors (e.g., environment, epigenetic) can contribute to the disorder.** Thus, instead of trying to predict the status of every input sample as affected case or unaffected control, as traditionally done in most classification approaches, *we propose to only predict a subset of samples as affected cases with very low false positive rate (FPR)*.

### Ultra-Accurate Disorder Prediction (UADP) problem

A positive diagnosis/prediction of a complex disorder (e.g., ASD) can have a severe negative psychological and economical impact on affected individuals and their family. For instance, a positive prediction of severe developmental disability during prenatal testing can result in a termination of pregnancy. *Thus, one of the main practical constraints in developing models and methods for prediction of a severe complex disorder is to guarantee a false positive rate (FPR) of virtually zero*. In other words, it is highly desirable not to have a false prediction of an input unaffected control as an affected case (similar assumption was made in other types of predictions [44]). Note, the UADP problem is different from traditional binary classification problems where *each sample* is assigned to one of the two classes (i.e., affected case or unaffected control). In the UADP problem, the goal is to predict a subset of samples as affected cases while all other samples are not assigned to any class. Note that intuitively this general idea behind UADP problem has been quite successful in handful of cases in practice, and has been the bases for screening some known genes with high penetrance for a specific disease. For example screening for mutations in *BRCA1/2* genes for breast cancer, or screening for LGD variants in few well-known genes with high penetrance for neurodevelopmental disorders (e.g. screen of FMR1 or MECP2 for fragile-X or Rett syndrome). Another example is the screening of CNVs such as 16p11.2 deletion and duplications which can explain up to 1% of autism with low false positive rate. In all of these examples the tests will cover only a fraction of the disease of interest (like breast cancer, autism or intellectual disability) but guaranty a low false positive rate for prediction of the disease. In the UADP problem, we are formalizing this approach as a formal computational problem and extending to be applicable in theory to any disorder.

In this paper, we study the UADP problem and provide a framework for solving this problem using rare coding variants. We aim to develop a computational method for positive prediction of a significant fraction of affected cases (due to the prediction limitation from coding variants) with virtually zero false positive prediction of unaffected controls (due to the negative effects of false positive prediction). We choose autism and related disorders as a case study since we can utilize a rich dataset of *de novo* mutations. In addition, we also integrate the functional relationship to increase the prediction capability. Approaches such as the one presented in this paper are needed to translate the biomedical discoveries into actionable items by clinicians.

## 2 Results

### 2.1 Data Summary

We tested Odin for accurate prediction of neurodevelopmental disorders using the likely gene disruptive (**LGD**) *de novo* variants from WES and targeted sequenced samples with ASD or ID. Table 1 shows the total number of samples and LGD variants reported from the union of several publications on over 6,000 ASD/ID probands. The union dataset of *de novo* variants used from these publications can be found in [45].

**Table 1:** The total number of ASD/ID-affected probands (cases) and unaffected siblings (controls) used in this study.

| Class - ASD diagnosis<br>(affected or unaffected) | Study | Number of<br>samples | Number of<br>LGD variants | References |
| --- | --- | --- | --- | --- |
| **Affected ASD/ID probands** | Simons Simplex Collection (SSC) | 2,508 | 492 | [17, 29, 46–48] |
|  | Autism Sequencing Consortium (ASC) | 2,270 | 185 | [18] |
|  | Other studies | 1,329 | 74 | [49–51] |
|  | **Total cases** | 6,107 | 751 |  |
| **Unaffected siblings or controls** | Simons Simplex Collection (SSC) | 1,909 | 248 | [17, 29] |
|  | GoNL | 250 | 7 | [52] |
|  | Other studies | 208 | 11 | [19] |
|  | **Total control** | 2,367 | 266 |  |

For building the gene similarity matrix *P*, which is used to convert the input variant vector for every sample (i.e, **z_i_** = **x_i_** *× P*), we have used the combination of coexpression values between two genes during brain development [38] and the difference between the likelihood of observing LGD variants in general population [53]. We can trivially extend the matrix *P* to include additional data such as tissue specific networks [54]. We observed that using such a matrix to map the variant vector for each sample into new space results in significantly reducing the *l*_1_ distance of probands with each other (*p* < 1.6*e -* 16). This indicates that using such a transformation indeed helps in increasing the prediction power.

Considering the samples in Table 1, there are few genes with significant recurrence of *de novo* LGD variants in affected cases while having no *de novo* variant in unaffected controls. Any prediction model/method for ASD can be trivially extended to predict a sample as an affected case if they have an LGD *de novo* variant in any of these genes. Thus, in our test data we will not consider any samples with *de novo* variants in any of these genes that are recurrently mutated in our training data. We will call these samples trivial cases/samples and the remaining samples as nontrivial cases/samples. Note that there are nine genes with four or more LGD variants in union of these ASD/ID samples (Table 1) with no LGD variant in unaffected siblings and controls. These nine genes are *ADNP, ANK2, ARID1B, CHD2, CHD8, DSCAM, DYRK1A, SCN2A* and *SYNGAP1* and any sample with an LGD variant in any of these genes is considered a trivial case to predict and it is not considered in our analysis.

#### Naïve approach of utilizing predicted ASD/ID genes for solving UADP problem

We would like to emphasize that based on our analysis (of samples in table 1) the approaches such as using the ASD/ID gene rankings in recently published methods (e.g., [39]), or selecting genes based on intolerance to LGD variant and expression in the fetal cortex **does not provide an acceptable solution to the UADP problem**. For example considering any sample with LGD variant in the top 100 genes from [39] as an affected case will have a false positive rate (FPR) of > 1% and true positive rate (TPR) of < 2.5%. Similarly predicting any samples with LGD variant in top genes based on the intolerance to LGD variant (from ExAC data) and high expression in cortex region during early fetal development (from CSEA tool http://genetics.wustl.edu/jdlab/csea-tool-2/) as an affected case will have a false positive rate of > 1% and true positive rate of < 2.8%.

### 2.2 Unicolor clustering with dimension reduction

We will first show that the proposed iterative method in Section 4.2.3 for solving the WUCDR problem (number of dimensions selected < 10) does in fact greatly improve the number of cases covered in comparison to the unweighted result (considering all dimensions with weights *w*_*i*_ = 1). As shown in Table 2, the optimal result found using the input was only able to cover 45 cases (only 24 of the cases not having LGD variants in recurrently mutated genes, i.e., the number of nontrivial cases covered). However, the WUCDR approach **converges** in less than five iterations and was able to cover over 71 cases (40 of the cases not having LGD variants in significantly recurrent mutated genes). Thus, our iterative approach for WUCDR improves the number of affected cases covered by over 60% using less than 10 dimensions. We also investigated the “density” of cases inside each selected region. The density was defined as the ratio between the number of affected cases covered and the radius *r*. We observed that not only the number of cases covered was improved using the WUCDR approach but also the density was increased (see Table 2) per iteration.

**Table 2:** Number of ASD/ID affected probands from training dataset covered in each iteration.

| Iteration | Number of cases covered | Number of non-trivial cases covered | Density (case/radius) | Number of dimensions (before - after) rounding |
| --- | --- | --- | --- | --- |
| 0 | 45 | 24 | 0.11 | $\geq 20,000$ |
| 1 | 53 | 32 | 138.41 | (15 - 9) |
| 2 | 66 | 39 | 176.64 | (7 - 7) |
| 3 | 70 | 39 | 179.92 | (10 - 9) |
| 4 | 71 | 40 | 185.49 | (10 - 9) |

### 2.3 ASD/ID disease prediction results

We compared the Odin framework in predicting affected ASD cases in comparison to different classification methods. We have used the k-NN classifier (various k-values 1 ≤ *k* ≤ 20), support vector machines (SVM) [55], and (lasso and elastic-net) regularization of generalized linear models [56]. We are specifically interested in comparing these methods in predicting ASD in *nontrivial* cases. We used the leave-one-out (LOO) technique to compare the Odin framework versus prediction power of k-NN, SVM classifiers, (lasso and elastic-net) generalized linear models. The exact commands used for running the k-NN, SVM and Glmnet for these experiments is provided in supplementary section 1.5. As our stated goal is to keep the false positive prediction of unaffected samples as cases close to zero, we will only consider the most conservative results for each method (false positive rate < 0.01). Odin’s true positive rate for predicting ASD is at least two times higher than the best k-NN result (for different values of k) and significantly higher than SVM for false positive rate < 0.01 (Figure 1). It is also significantly higher for different regularized generalized linear models (lasso and elastic net) for different input parameters of *α* (Glmnet implementation). For each of these tools, we used their intrinsic properties to control/limit the false positive rate for calculating the true positive rate. In k-NN we used the difference of number of affected cases and unaffected controls in the *k* closest neighbor; for SVM and generalized linear models we used the predicted probability (or distance) given by the libSVM [55] or Glmnet [56]. For both Glmnet and libSVM optimal set of training parameters were first picked considering all the data as input. That is the parameters of “gamma” and “cost” for libSVM and the parameter *s* for glmnet was set to optimal values learned using the full dataset (see supplementary section 1.5). For Odin, the weighted *l*_1_ distance of the sample to the selected center was used. Using these values we calculated the highest true positive rate for each method given the upper bound on the false positive rate value using LOO approach. Note that in Odin the full set of samples predicted as affected cases will have a false positive rate of less than 0.01.

**Figure 1:**
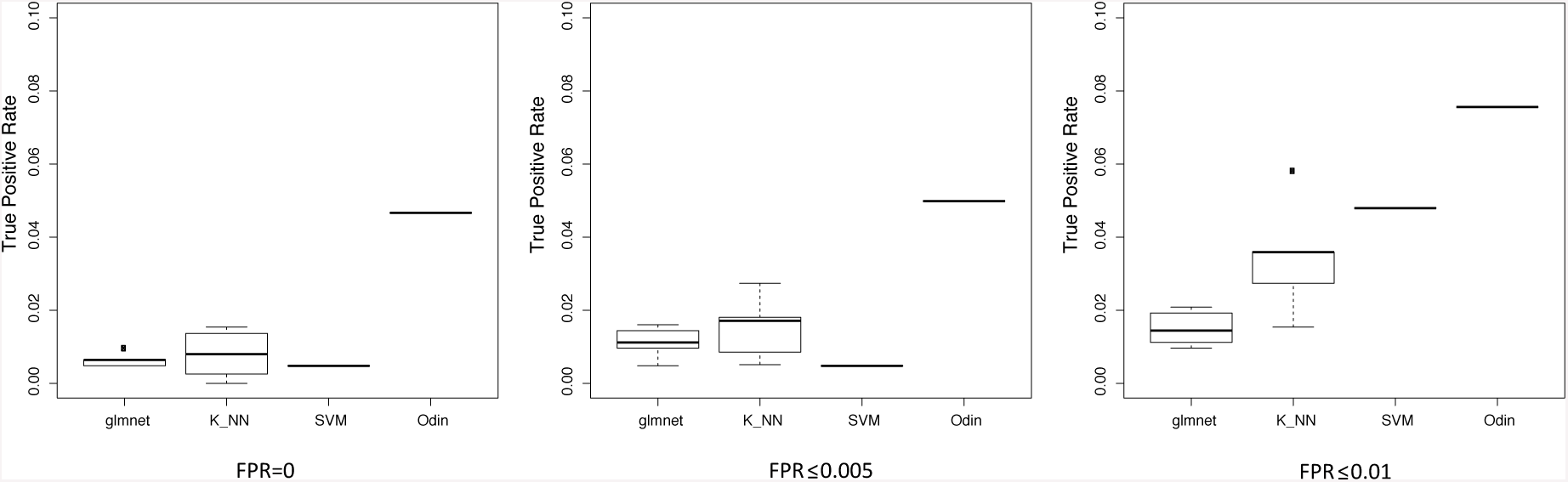
Prediction results on ASD/ID data set. For k-NN the boxplot shows the result of different values of k tested (1 ≤ *k* ≤ 20) and for GLmnet the boxplot shows the results for different values of *α* ∈ {0, 0.25, 0.5, 0.75, 1}.

Separately, we also compared Odin’s approach for prediction of ASD/ID against SVM for very low false positive rates by splitting the data into 20% and 80% representing the test data and train data respectively. We then applied Odin and libSVM for predicting the ASD cases on the testing set. We used 5-fold cross validation to select the best set of parameters for each method (e.g. “gamma” and “cost” for libSVM). As number of total input data with LGD variants is in general low and thus there is a large variance in performance of the methods based on different partitioning we repeated the above procedure 10 (independent) times to calculate average performance (FPR and TPR) of each method. For each of these runs we considered the same number of top score predictions by SVM as predicted by Odin to be cases. We then calculate the average of the FPR and TPR. In these set of experiments the average FPR for Odin predictions was 0.007 and the TPR was 0.11, while the FPR for SVM predictions was 0.012 and the TPR is 0.10. More interestingly, we noticed in these results that most of the true positive calls by SVM are from the samples with *de novo* variant in recurrently mutated genes (i.e. trivial cases). However, Odin was able correctly predict disease status for larger fraction of cases without *de novo* variants in recurrently mutated genes (non-trivial cases). For non-trivial cases the average TPR for Odin was 0.069 while the average TPR for SVM was 0.032.

We also showed Odin outperforms other approaches in predicting developmental delay disorder (DDD) using the *de novo* LGD variants based on the samples in deciphering developmental delay study [57] (please see the supplementary material section 1.2 for the developmental delay disorder analysis).

### 2.4 Autism and related disorders gene prediction and ranking

Odin is a framework to predict with ultra-accuracy a subset of samples that will develop ASD given the *de novo* variation; however, it can also be used to predict some novel ASD genes. We have utilized Odin to rank all genes for the potential impact of a *de novo* LGD variant disrupting them. Note that similar to the predictions of ASD/ID for subsets of samples made by Odin, if a gene is not selected does not mean it is not an ASD/ID gene. We used the ASD and Siblings variants (Table 1) as the training data and calculated the weighted *l*_1_ distance to the selected center. Ranking all the genes based of the calculated distance, we clearly see an enrichment of known ASD genes closer to the center (Figure 2a). For the set of known ASD genes we used the union of SFARI high-confidence genes and known syndromic genes.

**Figure 2:**
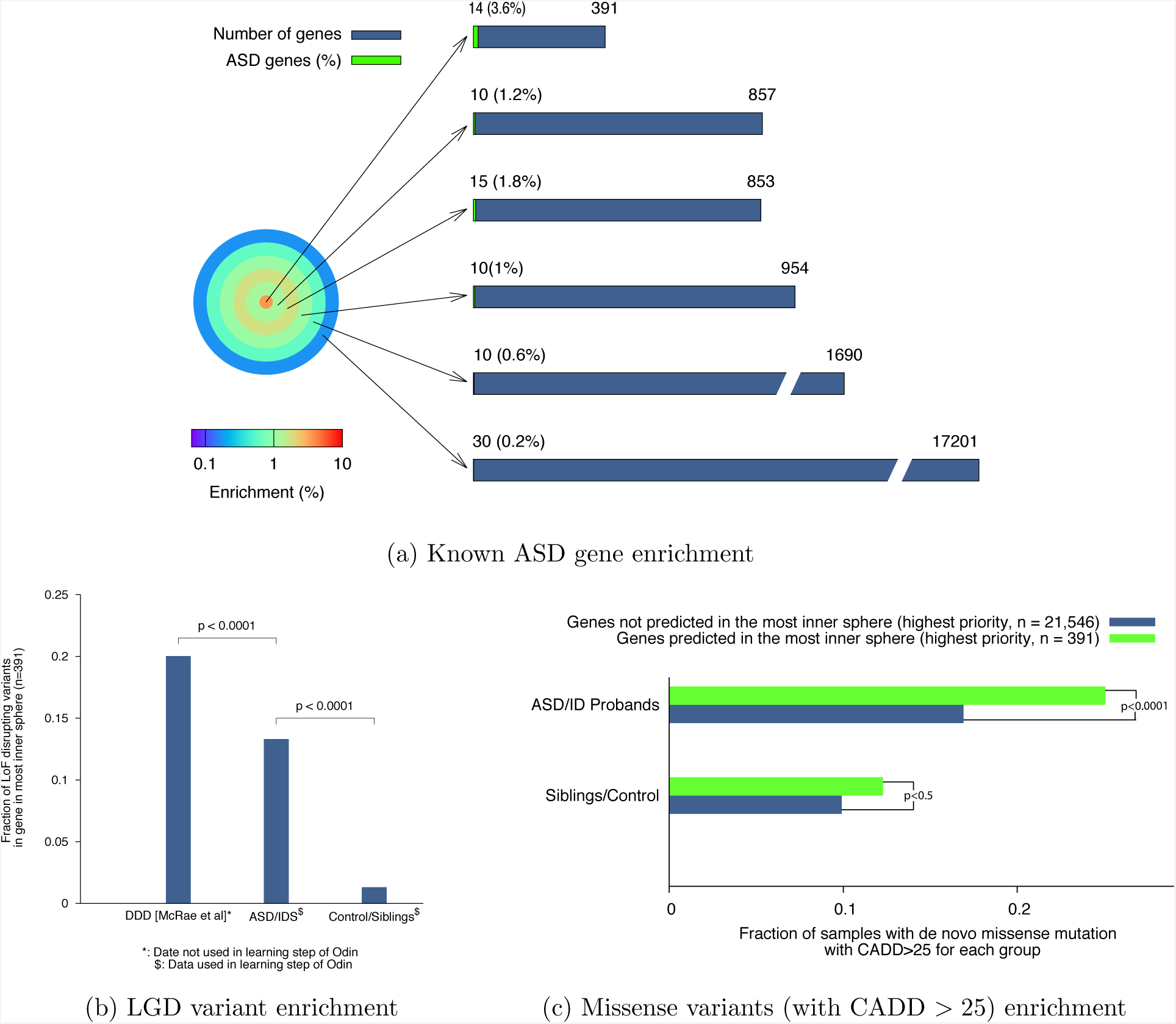
a) The set of genes which are closer - based on the weighted *l*_1_ distance - to the center selected by Odin are more enrichment in known ASD genes. b) Significant enrichment of LGD variants in ASD/ID/DDD disrupting the genes in inner most sphere (391 total genes). c) Significant enrichment of severe missense variants (CADD> 25) disrupting these 391 genes in ASD/ID/DDD probands.

Our analysis also indicated that there are 391 genes where an LGD variant on them will result in a sample falling inside the predicted area of interest (i.e., the inner most sphere/circle in Figure 2a). These 391 genes indicate a set of genes in which Odin has predicted with high probability that their disruption will cause significant (neuro)developmental disorder. Furthermore, there was significant enrichment of LGD variants in these 391 genes in the DDD set (which was not used in the training) versus the ASD set (which was used in the training) as shown in Figure 2b. Interestingly, this clearly indicates that even after normalizing based on expected LGD variants for each disease group the more severe samples tend to be more enriched in LGD variants than their less severe autism samples disrupting these selected 391 genes (Figure 2b). There is also an enrichment of severe *de novo* missense variants (i.e., with CADD score > 25) disrupting these genes (Figure 2c) in affected autism/ID/DDD probands while no such enrichment is seen for control/sibling samples (Figure 2c). Furthermore, we used the *de novo* variants reported in 520 whole-genome sequenced (WGS) samples [58] which were void of LGD variants to also investigate the *de novo* variants disrupting the **non-coding regulatory regions of these genes** (see supplementary section 1.3). We observed that the non-coding regulatory elements of these 391 genes are significantly disrupted by *de novo* variants in probands versus siblings (*p* < 0.004 - Supplementary Figure 3).

In the Simons Simplex Collection (SSC) we also observed not only that probands with LGD variant tend to have a lower IQ than probands without *de novo* LGD variants, but also the probands with *de novo* LGD variant disrupting one of the genes in the most inner sphere (the 391 genes) have lower IQ than other probands with *de novo* LGD variants (Figure 3a). It is been known that their is a large male to female bias in autism (estimate to be over 4 to 1). Note that in the Simons Simplex Collection (SSC) there are a total of 2478 male probands and 396 female probands (over 6:1 ratio). However, the difference between the number of samples with LGD *de novo* variants in the selected 391 genes in the inner most sphere is 31 to 16 (around 2:1 ratio). This indicates that there is much smaller gap for sex difference for ASD samples with *de novo* LGD variants in the predicted ASD genes by Odin (Figure 3b). We were also interested to see if there are any specific enrichment of expression of these top 391 genes selected in human brain. We used the online tool CSEA [59] (http://genetics.wustl.edu/jdlab/csea-tool-2/) to study the expression profile of these 391 genes. Interestingly, the only significant expression we observed was on the early fetal development and mid-early fetal development of brain (Figure 3c). No significant expression of these genes in any tissues in adult human or mouse brain was observed.

**Figure 3:**
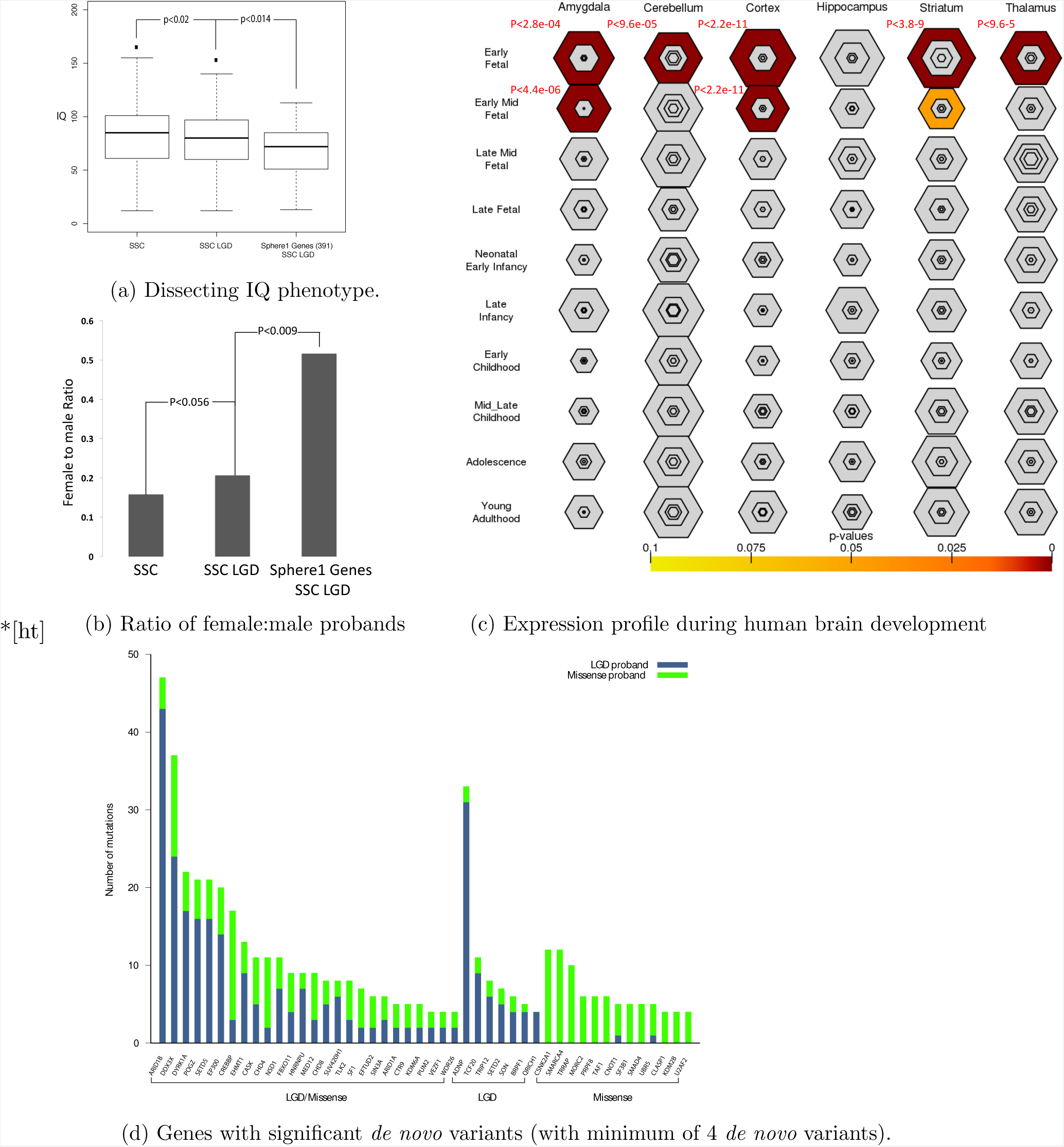
Properties of genes selected by Odin in the inner most sphere (391 total genes). a) The ASD probands with LGD variant in these genes have significant lower IQ than other ASD probands with LGD variant. b) The gap between female:male ratio is significantly smaller for probands which have LGD variants in these 391 genes. c) These genes are expressed during early fetal and early mid-fetal brain development. The reported p-values are calculated by the CSEA tool (http://genetics.wustl.edu/jdlab/csea-tool-2/) after Benjamini-Hochberg statistical correction. d) The subset of genes in inner most sphere which are significantly enriched in variants in probands.

Finally, we could group these 391 genes based on observing significant *de novo* LGD and/or missense variants in affected probands (Figure 3d). We utilized the predicted probability of observing a missense or LGD *de novo* variant per sample for each gene [46] to calculate the p-values of observed *de novo* variants in the affected samples. The set of genes with only significant missense *de novo* variants observed in cases potentially indicates genes in which an LGD variant will be incompatible with life (i.e., essential genes). However, a missense mutation can result in a severe (neuro)developmental disorder. These genes include *CSNK2A1, SMARCA4, TRRAP, MORC2, PRPF8, TAF1, CNOT1, SF3B1, SMAD4, UBR5, CLASP1, KDM2B,* and *U2AF2*. The list of these 391 genes is provides as supplementary table 1.

### 2.5 Pathways

We were interested in studying the properties of the samples that Odin correctly predicted as an affected case. We used the tool David (v.6.7) [60] for the discovery of enriched GO-terms and KEGG pathways for the genes mutated in these samples. We found that for the ASD/ID samples in table 1, Odin was able to correctly predict the ASD status of the samples that have *de novo* variants in genes in the Wnt pathway or chromatin regulators (Figure 4a). Similarly, correctly predicted development delay disorder (DDD) study samples [57] had mutations in chromatin modification and transcription regulation genes (Figure 4b). Note that these pathways were previously indicated to be a major contributing factors in neurodevelopmental disorders [18, 37, 38, 61, 62].

**Figure 4:**
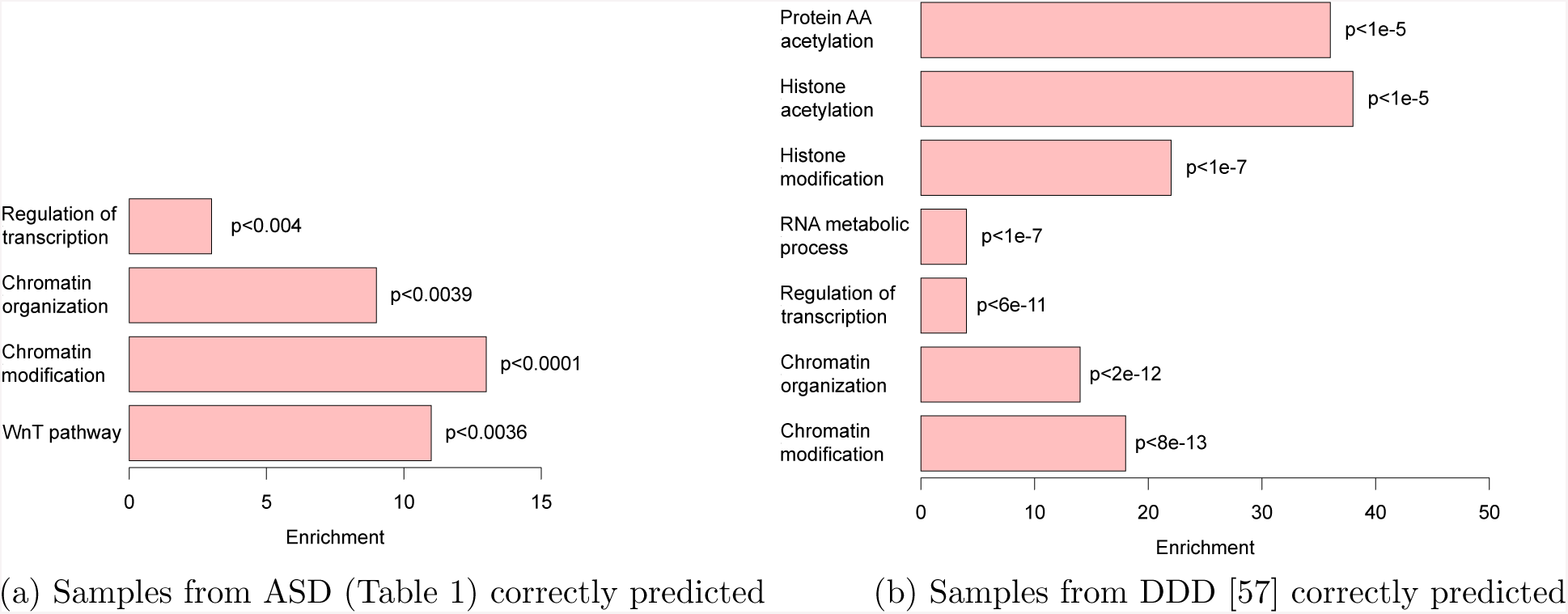
Pathways and Gene Ontology (GO) enrichment.

## 3 Discussion

In this paper, we have formalized a computational problem for solving a specific version of complex disorder prediction that enforces a virtually zero false positive prediction as to make it directly applicable for diagnosis in clinical settings. We denoted this specific problem as Ultra-Accurate Disorder Prediction (UADP) problem, and have shown that simple approaches of utilizing the predicted ASD/ID gene rankings is not a viable solution for UADP problem. We also introduce the framework Odin for solving the UADP problem in autism and related disorders using *de novo* LGD variants. Our evaluation of the experimental data shows an advantage of Odin in predicting ASD/ID using LGD *de novo* genetic variants. The proposed framework can be extended to take into account not only LGD mutations but also *missense mutations* to increase the power of the model in predicting a higher percentage of affected cases. As we have shown, there is clear enrichment of severe missense mutations (CADD score > 25) to genes closer to the predicted center. We can adapt evolutionary based scores (e.g., CADD score [63] or polyphen-2 score [64]) to define an additive summarization function to assign a disruption score for each gene (i.e., a continuous value in comparison to a binary value as done in this paper). In addition, we can integrate other information, such as protein interaction [65, 66], tissue-specific networks [54] or the regulation of specifically related pathways such as Wnt [62] or mTOR [67], to increase the prediction capability. For the algorithm, we can improve the first guessed solution (in the first step) of WUCDR (see Section 4.2.3) by utilizing algorithmic techniques in geometry. Our proposed framework here can be extended for predicting the risk of other neurological disorders, such as schizophrenia, epilepsy or Alzheimer.

Note that Odin is not meant to replace other approaches for disorder gene discovery and ranking, but rather for accurate prediction of the disorder in a subset of cases given the genetic variation. One of the drawbacks of this approach is that it only can work if the penetrance of the genetic variation to cause the disorder is very high. It is not applicable in cases which the goal is to only predict if the probability of disorder is significantly higher than the general population.

Finally, the pathways indicated by Odin (i.e. WnT pathway and chromatin remodelers) are some of the main pathways and modules found to be important for autism and related disorders [37, 38]. However, there are other well-known pathways which are also contributing to the disorder which have not been indicated by Odin (e.g., long-term potentiation and synaptic function). The main reason is that the current formulation of Odin only considers one center and thus genes with similar functionality to one module/pathway are mainly covered and other modules are not being covered. We believe an extension of Odin which considers multiple centers can significantly improve the results and pathways covered.

## 4 Methods

### 4.1 UADP problem definition and notations

In the UADP problem we are trying to maximize the number of samples correctly predicted as affected case, while the number of unaffected controls falsely predicted as affected cases must be extremely small. Another way to look at this problem is that we are trying to select a subset of samples such that the total number of unaffected controls picked is negligible while the number of affected cases selected is maximized. Finally, note that because of low recurrence of the same rare and *de novo* variants, we will be using a summarization of coding rare variants based on their effect on each gene and the biological function disrupted to increase our power for prediction.

#### Training Data

Let *n* and *m* be the number of genes and the total number of samples respectively. The *LGD (likely gene disruptive) mutation profile* of the *i*^*th*^ sample is a binary row vector 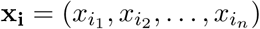 where

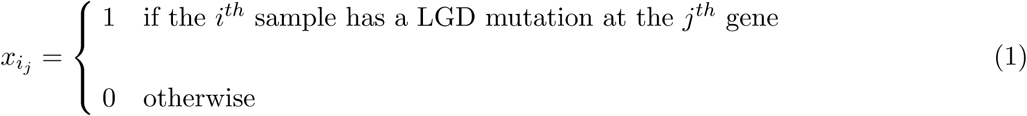

An assumption here is that an LGD mutation will completely knockout or disrupt the copy of the affected gene in the sample. The *diagnosis result (or class)* of the *i*^*th*^ sample is a binary value *y*_*i*_ where

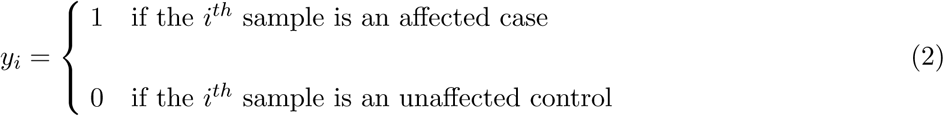

A *dataset D* of *m* input samples is a set of *m* pairs *D* = {(**x_1_**, *y*_1_), (**x_2_**, *y*_2_), *…,* (**x_m_**, *y*_*m*_)} where each pair (**x_i_**, *y*_*i*_) represents the LGD mutation profile and the diagnosis result respectively of the *i*^*th*^ sample. We define the unaffected control set and the affected case set as *D*_*control*_ = {**x_i_***|*(**x_i_**, *y*_*i*_) ∈ *D, y*_*i*_ = 0} and *D*_*case*_ = {**x_i_***|*(**x_i_**, *y*_*i*_) ∈ *D, y*_*i*_ = 1}, respectively.

#### Gene similarity score

We will use the functional similarity between genes to increase the statistical power in disorder prediction. The assumption is that disruption of genes with similar functionality will result in similar phenotypes. Thus, we would like to develop a framework that can include the similarities between genes (mutational landscape and function) as an additional signal for disease prediction. We denote such a matrix by *P* ∈ [0, 1]^*n×n*^ where *P*_*i,j*_ indicates the similarity between genes *i*^*th*^ and *j*^*th*^ and potentially how disruption of one gene can affect the other gene. For neurodevelopmental disorders such as ASD our goal is to build the matrix *P* to reflect functional similarity of genes during brain development. One way to calculate such a score for any pair of genes is based on using the coexpression of genes during brain development. Coexpression between two genes *i* and *j* is denoted by *R*(*i, j*) and is calculated to represent the expression similarity of these two genes in different conditions and tissues. Similar to previous practices, we are using the Pearson correlation of expression profiles between two genes in different conditions as the coexpression value [38, 68–71]. Coexpression has been shown to be a powerful indicator of functional similarity of two genes for neurodevelopment. We also include the similarity of likelihood of observing LGD mutation (pLI) between the two genes in the population [53] for building this matrix. Assuming we are given multiple matrices capturing the similarity of genes with each matrix using different biological concepts, we will use the minimum of similarity scores of two genes among different matrices to build matrix *P*. In another words, if we have two matrices of gene similarity *P* ^*′*^ and *P* ^*″*^, the matrix *P* is built by assigning 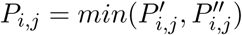. Of course, the choice of these matrices and the way we combine them together can be changed without any need to change the underlying framework and proposed methods. The details of different datasets used to build the matrix *P* for this study is provided in Section 2.1. We will convert every sample by multiplying the vector **x_i_** by matrix *P* to produce new vectors **z_i_** = **x_i_** *× P*. We will denote the set of samples *D*_*control*_ and *D*_*cases*_ converted by the gene similarity matrix *P* as *D*′_*control*_ = {**z_i_** = **x_i_** *× P |* **x_i_** ∈ *D*_*control*_} and *D*^*′*^_*case*_ = {**z_i_** = **x_i_** *× P |* **x_i_** ∈ *D*_*case*_}.

### 4.2 Odin framework

In this subsection, we will introduce the intuition behind our framework Odin (**O**racle for **DI**sorder predictio**N**) as a practical solution for the UADP problem. To build such a conservative prediction model, Odin will intuitively predict an input/test sample to be an affected case if and only if it satisfies two conditions:

1. The input sample is “close” to many affected case samples
2. The input sample is “far” from any unaffected control sample

For satisfying the first condition, we simply use the nearest neighbor approach using a distance function (e.g., Euclidean distance). The closest neighbor of the input sample among the training data should be an affected case so that input sample passes the first condition.

For satisfying the second condition, we will initially develop a novel algorithm that first finds a region (after dimension reduction) containing a significant number of affected cases and does not contain any unaffected controls. This cluster is denoted as *unicolor cluster*, as it only includes the affected cases. The input sample passes the second condition if it falls inside of this unicolor cluster. We denote the problem of finding such a cluster as Unicolor Clustering with Dimensionality Reduction (UCDR). We prove that this problem is a NP complete problem (section 1 in the supplemental material) and can not be solved efficiently. Therefore, we propose a relaxation of UCDR that we denote as Weighted Unicolor Clustering with Dimensionality Reduction (WUCDR). In the remainder of this section, we will first formalize the UCDR and WUCDR problems and then present an iterative algorithm to solve the WUCDR problem.

#### 4.2.1 Unicolor Clustering with Dimensionality Reduction (UCDR) problem

In the UCDR problem we have a set of *red* and *blue* points in *n*-dimension space ℝ^*n*^ representing unaffected controls (i.e., *D*^*′*^_*control*_) and affected cases (i.e., *D*^*′*^_*case*_), respectively. Furthermore, we have an upper bound on the number of dimensions to consider (dimension reduction/feature selection) denoted by *k*. The goal of the UCDR problem is to discover a subset of dimensions with cardinality *k* (*k* ≪ *n*), a center point **c** ∈ ℝ|k| and a constant *r* such that after mapping all the blue and red points to the reduced *k* dimensions the following objective and constraint hold:

- **Objective:** *maximize* the total number of blue points with “distance” less than *r* to center **c**.
- **Constraint:** there is no red point with “distance” less than *r* to center **c**

As a general rule any metric distance function (e.g., Euclidean distance) can be used for the UCDR problem. However, we are using the *ℓ*_1_ distance since it is concordant with the additive model used in common variant studies. The *ℓ*_1_ distance between two points (*a*_1_, *a*_2_, *…, a*_*n*_) and (*b*_1_, *b*_2_, *…, b*_*n*_) is defined as 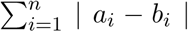. We will denote the region contained with distance *r* from center **c** ∈ ℝ*|k|* as *area of interest 𝒜*(**c**, *r*). Furthermore, any affected case **z_i_** ∈ *D*^*′*^_*case*_ inside the area of interest (i.e., *ℓ*_1_ (**c**, **z_i_**) ≤ *r*) is considered covered by this area.

Note that the intuition behind the dimension reduction is to avoid the overfitting issue raised as a result of a large number of dimensions (> 20, 000 genes) and a small number of training samples. In practice, we will require that the number of selected dimensions be less than *O*(log_2_(*m*)) (i.e., *k* = *O*(log_2_(*m*))) where *m* is the total number of training samples (both cases and controls).

#### 4.2.2 Weighted Unicolor Clustering with Dimensionality Reduction (WUCDR) Problem

Since UCDR problem is NP-complete (see Section 1 in the supplemental material), we will define a relaxation, where we assign (continuous value) weights to the dimensions. We denote this problem as the Weighted Unicolor Clustering with Dimensionality Reduction (WUCDR) problem. More formally, in addition to selecting *k* genes/dimensions, we also have to assign weights 0 ≤ *w*_*i*_ ≤ 1 to each gene/dimension *i* and use the **weighted** *ℓ*_1_ as the distance metric for clustering (we use the notation *w ℓ*_1_ to represent weighted *ℓ*_1_). In the rest of the paper, we will define the weighted *ℓ*_1_ distance function between two input points **a** and **b** with weights **w** (in *n* dimensions) as 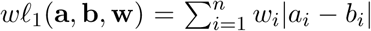. Note that as we are only allowed to select *k* dimensions thus, over *n - k* other dimensions will have weight zero.

#### 4.2.3 Iterative solution for WUCDR

Here we propose an iterative approach consisting of two main steps to solve the WUCDR problem.

In the first step, given a set of weights **w**, we find the optimal center **c** and radius *r* to cover a maximum number of affected cases (blue points) in the area of interest *𝒜*(**c**, *r*) (note that the area of interest is considered using weighted *ℓ*_1_ distance). In the second step, we try to find a new set of weights **w** given the center **c** and the radius *r*.

#### First Step

Given the weights **w** = (*w*_1_, *w*_2_, *…, w*_*n*_) (all the weights are assigned to 1 at the first iteration), find a center **c** and constant *r* such that

1. all red points have a weighted *ℓ*_1_ distance greater than *r* to center **c** and
2. the number of blue points, which have weighted *ℓ*_1_ distance less than *r* to center **c**, is maximized.

In general, finding such a center is a hard problem in *n* dimensional space and can be very time-consuming. Thus, we will relax the problem only to consider the blue points as a potential center **c**. This can be done trivially in polynomial time by considering every blue point as potential center and picking the optimal one. Given a center **c**, radius *r* and the weights **w** we can easily calculate the affected cases (i.e., blue points) covered by the area of interest. Let set *S* denote the covered (blue) points (i.e., affected cases), which will be used in the next step for updating the weights.

#### Second Step

Given a center **c** and the set of blue points *S*, covered by the area of interest found in the first step, we will calculate new weights **w** (for each dimension). The objective is to decrease the weighted *ℓ*_1_ distance of points in the set *S* to center **c**, while increasing the weighted *ℓ*_1_ distance of points in the set *S* to the red points (*D*^*′*^_*Control*_). We will solve the linear programming (LP) problem below to find these new weights

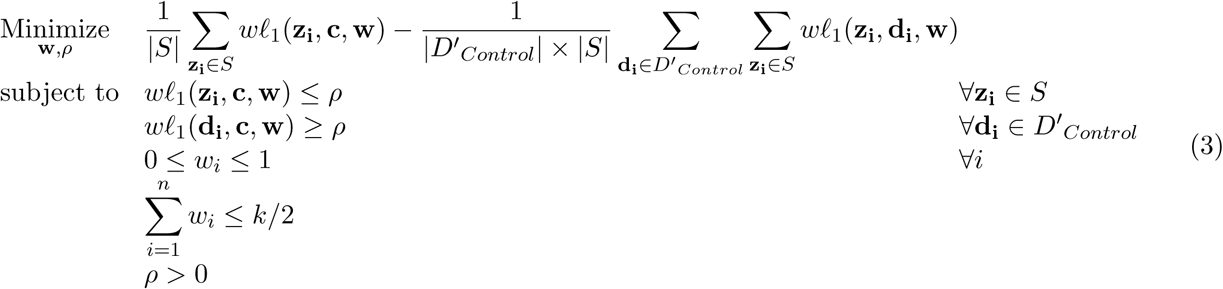

Note that in the above LP problem only **w** and *ρ* are unknown variables, while the set *S* and center **c** are calculated in the first step of the method. The constraints in the above LP problem will find a set of weights that are guaranteed to have all of the points in set *S* closer to the selected center **c** than any red point. Furthermore, these weights try to squeeze the (blue) points in *S* further closer to the center **c**, while increasing the distance of red points to the (blue) points in the set *S*. The objective function of the above LP problem has two main terms. The first term aims to reduce the average distance between points in the set *S* and the center **c**. Simply stated, the new weights **w** would try to make blue points covered in first step (i.e., point in set *S*) get closer to the center **c** (note that both **c** and *S* are from the previous step, *not* variables in this LP problem). The second term aims to increase the average weighted *ℓ*_1_ distance of all red points to the blue points in set *S*. Finally, among the weights produced we will only keep the top *k* weights and convert all of the remaining weights to 0. Note that because of the condition 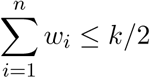, we are guarantied to be able to keep any dimension with value > 0.5 from the LP solution.

#### Odin framework using WUCDR

As mentioned in the Section 4.2, two conditions should be satisfied for a sample to be predicted as a potential affected case by Odin. The first condition is that the nearest neighbor to the samples should **not** be an unaffected control. Odin uses the *ℓ*_1_ distance function for finding the nearest neighbors of any test sample. The second condition is that the input sample should fall inside the *area of interest 𝒜*(**c**, *r*) after performing the same dimension reduction mapping using weights **w** (note that **c**, *r* and **w** are found by the iterative solution of WUCDR).

## 5 Acknowledgment

We would like to thank Evan E. Eichler, Tychele N. Turner, Phuong Dao, Farhad Hormozdiari, Madeleine Geisheker and Tonia Brown for reading the paper and providing comments.

## 1 Supplementary materials

### 1.1 Complexity of the UCDR problem

We show that an instance of the decision version of the UCDR problem is NP-complete.

#### Remark 1

Given a set of positive (rational) numbers. The problem of determining if there exists two disjoint nonempty subsets whose elements sum up to the same value is NP-complete [Woeginger, G. J., & Yu, Z. (1992). On the equal-subset-sum problem. Information Processing Letters, 42(6), 299-302].

The problem in Remark 1 was called *“equal subset sum problem”*. Notice that the pair of two subsets in the solution is not necessary a partition (i.e. there may be some elements that are in the original set but are not in either of these two sub-sets).

#### Theorem 2.

Given a set of points in a n-dimension space where each point was assigned a color either blue or red. The problem of determining if there exists a non-empty dimension subset and a center point such that all blue points are not farther to that center point in comparison to red points (by the L_1_ norm in the reduced dimension space) is NP-complete. We call the problem “UCDR decision problem”.

*Proof.* We will reduce the equal subset sum problem (Remark 1) to a special instance of the UCDR decision problem.

Assume we are given a set of positive rational numbers *A* = {*a*_1_, *a*_2_, *…, a*_*n*_}. We create two blue points *B*_1_ = (*a*_1_, *a*_2_, *…, a*_*n*_), *B*_2_ = (*-a*_1_, *-a*_2_, *…, −a*_*n*_) and one red point *R* = (0, 0, *…,* 0). We consider the UCDR decision problem of three points *B*_1_, *B*_2_ and *R*. Suppose that this UCDR decision problem has a solution that includes a dimension subset *I* = {*i*_1_, *i*_2_, *…, i*_*d*_} ⊆ {1, 2,…, n} and a center *C*.

Now we only consider the reduced space with *d* dimensions from *I*. We denote 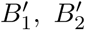, and *R*^*′*^ as the corresponding points of *B*_1_, *B*_2_, and *R* respectively in the reduced space.

Let *H* be the smallest (by volume) *L*_1_ norm ball that has the center *C* and contains both 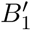 and 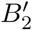. Thus 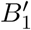 or 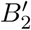 (or both) must be on a facet of *H*, we can assume 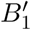 is on a facet of *H* without losing generality. Since *H* is convex and 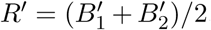, *H* also contains *R*^*′*^. But if 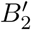 is not on the same facet of 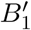, then *R*^*′*^ will be inside *H* and thus 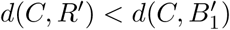. Therefore, both 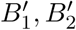 and *R*^*′*^ must be on the same facet of *H*. Let *F* be that facet, since *H* is a *L*_1_ norm ball then any point 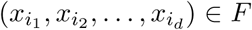 must satisfy an equation that has the form

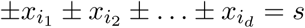

Since *R*^*′*^ = (0, 0,…, 0) ∈ *F*, so *s* must be 0. Thus we can re-write the equation as

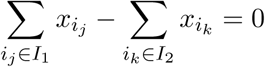

where *I*_1_ ∩ *I*_2_ = ∅ >and *I*_1_ *∪ I*_2_ = *I*. Since 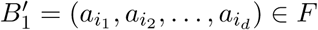 then

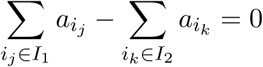

but both 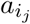 and 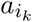 are in *A* that contains positive numbers only so *I*_1_ ≠ ∅ and *I*_2_ ≠ ∅. Therefore, the pair of two sets 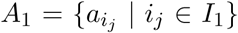 and 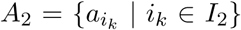 is a solution of the equal subset sum problem of the set *A*.

Thus, a solution of the UCDR decision problem is also a solution of the equal subset sum problem. Conversely, we can also easily verify that a solution of the equal subset sum problem is also a solution of the UCDR decision problem. Therefore, if we can solve the decision version of UCDR then we can solve the equal subset sum problem which is NP-complete (Remark 1). Since it is easy to verify this problem is in NP, it is also NP-complete.

### 1.2 Developmental delay disorder (DDD) prediction using Odin

In addition to the samples reported in Table 1, a set of over 4000 trios with developmental delay disorder (DDD) were whole-exome sequenced [57]. Since linear model (i.e. Glmnet) was not suitable for this prediction problem (as seen in Figure 1), we only compared Odin against the k-NN approach (1 ≤ *k* ≤ 10) using the dataset in Table 1 as a training set and the (nontrivial) DDD affected cases [57] for testing (Supplementary Figure 1). Note that we used the parameters learned from the previous set of experiments (Section 2.3) to control the false positive rate. The Odin method was able to accurately predict a higher fraction of nontrivial DDD probands in comparison to k-NN approach (Supplementary Figure 1a) using the ASD/ID samples as training. We further investigated the overlap between nontrivial DDD affected samples which were correctly predicted by Odin and 1-NN (nearest neighbor) approach (Supplementary Figures 1b).

**Supplementary Figure 1:**
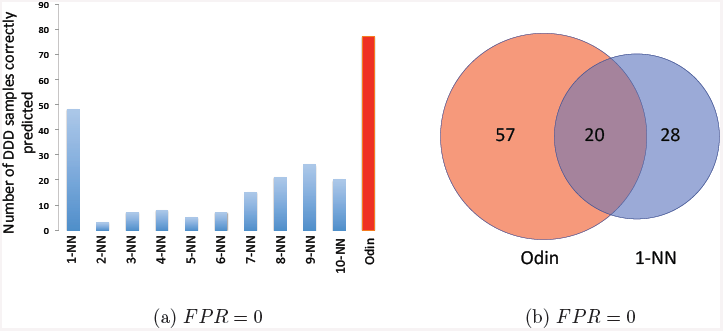
Prediction results on DDD data set

Interestingly, there are significant number of samples which were correctly predicted only by one of the methods, which indicates an approach which combines different methods can even outperform Odin. We further analysis Odin’s capability in accurate prediction of DDD probands with FPR < 0.01 while using the ASD/ID data (samples in Table 1) as training.

**Supplementary Figure 2:**
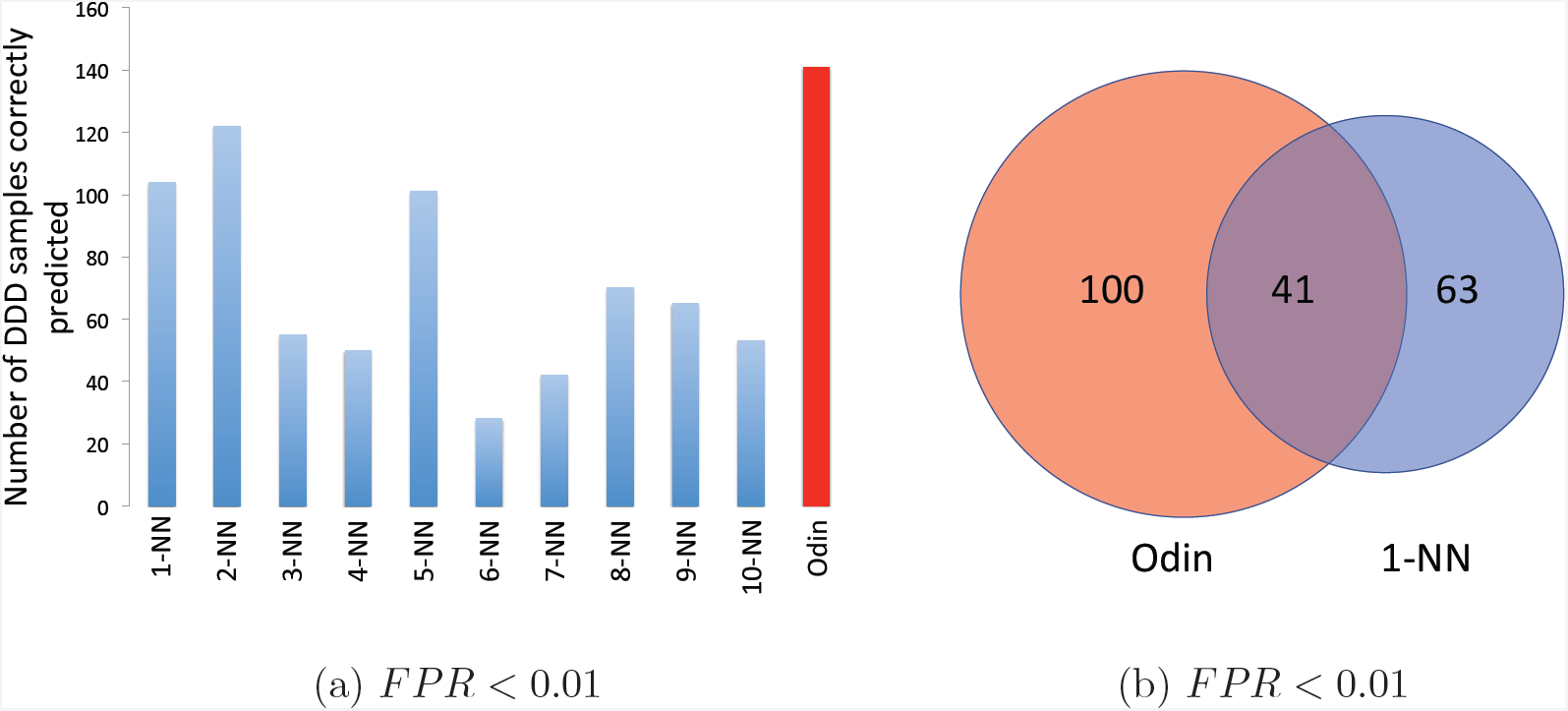
Prediction results on DDD data set

### 1.3 Enrichment of non-LGD variants in ASD probands disrupting the selected genes and their regulatory elements

A total of 516 ASD simplex families from SSC were recently WGS and *de novo* variants in the affected probands and unaffected sibling were predicted and validated [58]. Note that these families were selected *to be void of LGD variants based on whole-exome sequencing*. Thus, they were not part of the samples which contributed to Odin training. However, we did observe a significant number of the affected probands in comparison of unaffected siblings had non-LGD coding and non-coding *de novo* variants disrupting the coding or the regulatory elements of the genes in the inner most sphere (Supplementary Figure 3). The subset of genes in the selected 391 genes in the inner most sphere, which had a *de novo* variants disrupting their coding or regulatory elements in probands or siblings, is depicted in Supplementary Figure 4. Furthermore, we also observed the significant enrichment after removing the known SFARI high confidence and syndromic autism genes from the set of 391 genes considered.

**Supplementary Figure 3:**
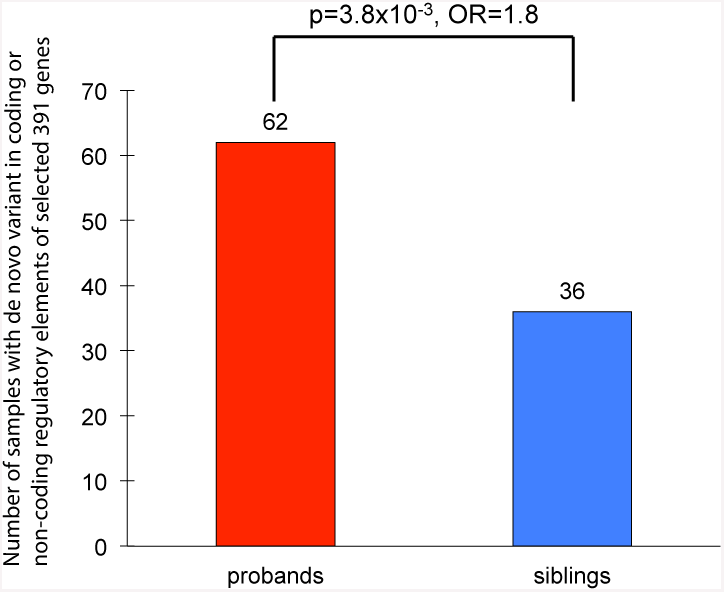
Enrichment of *de novo* non-LGD variants in WGS samples disrupting coding and regulatory regions of genes in inner most sphere (total 391) in affected probands versus unaffected siblings.

**Supplementary Figure 4:**
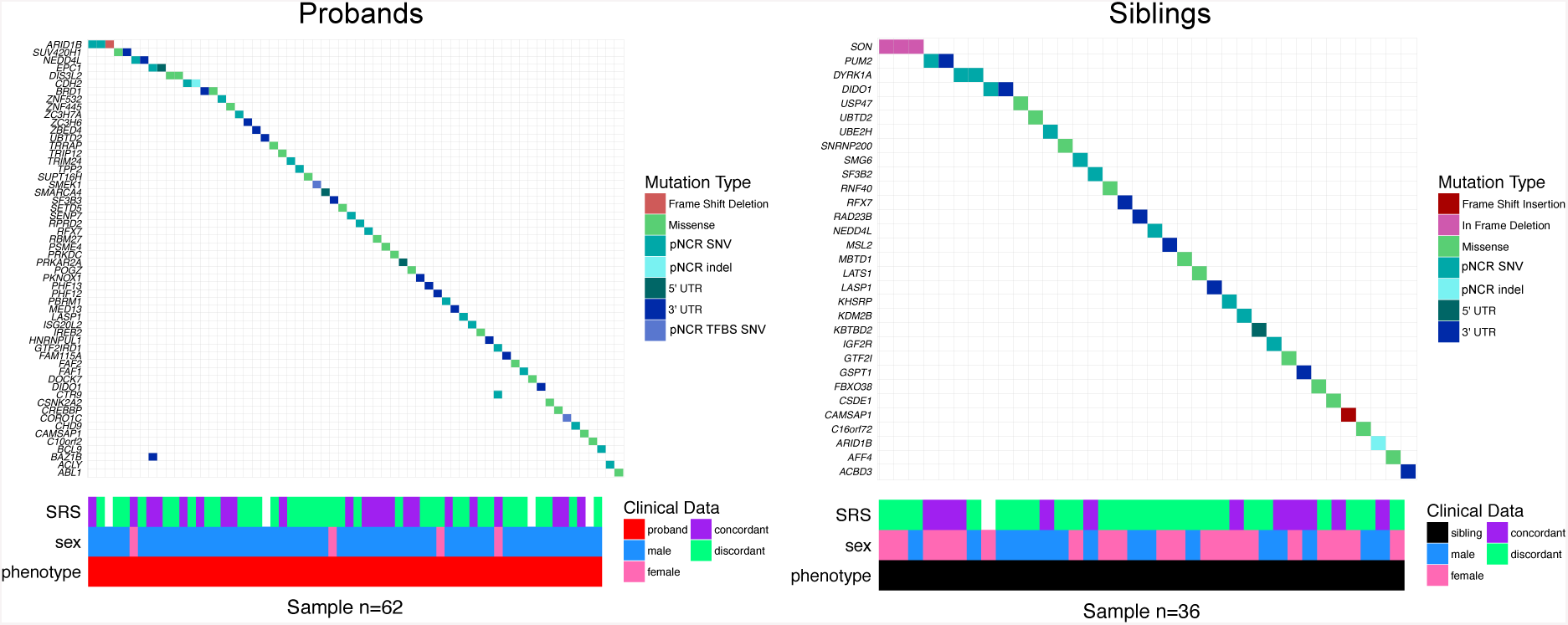
The non-LGD disruptive variants disrupting coding and regulatory regions of the genes in inner most sphere in ASD probands and siblings.

### 1.4 Protein interaction enrichment

We investigated the changes in genes degree in protein-interaction networks based on their weighted *ℓ*_1_ distance to the center found using Odin. There is an interesting correlation between distance calculated by Odin for each gene and the average degree of that genes in protein-interaction networks (Supplementary Figure 5).

**Supplementary Figure 5:**
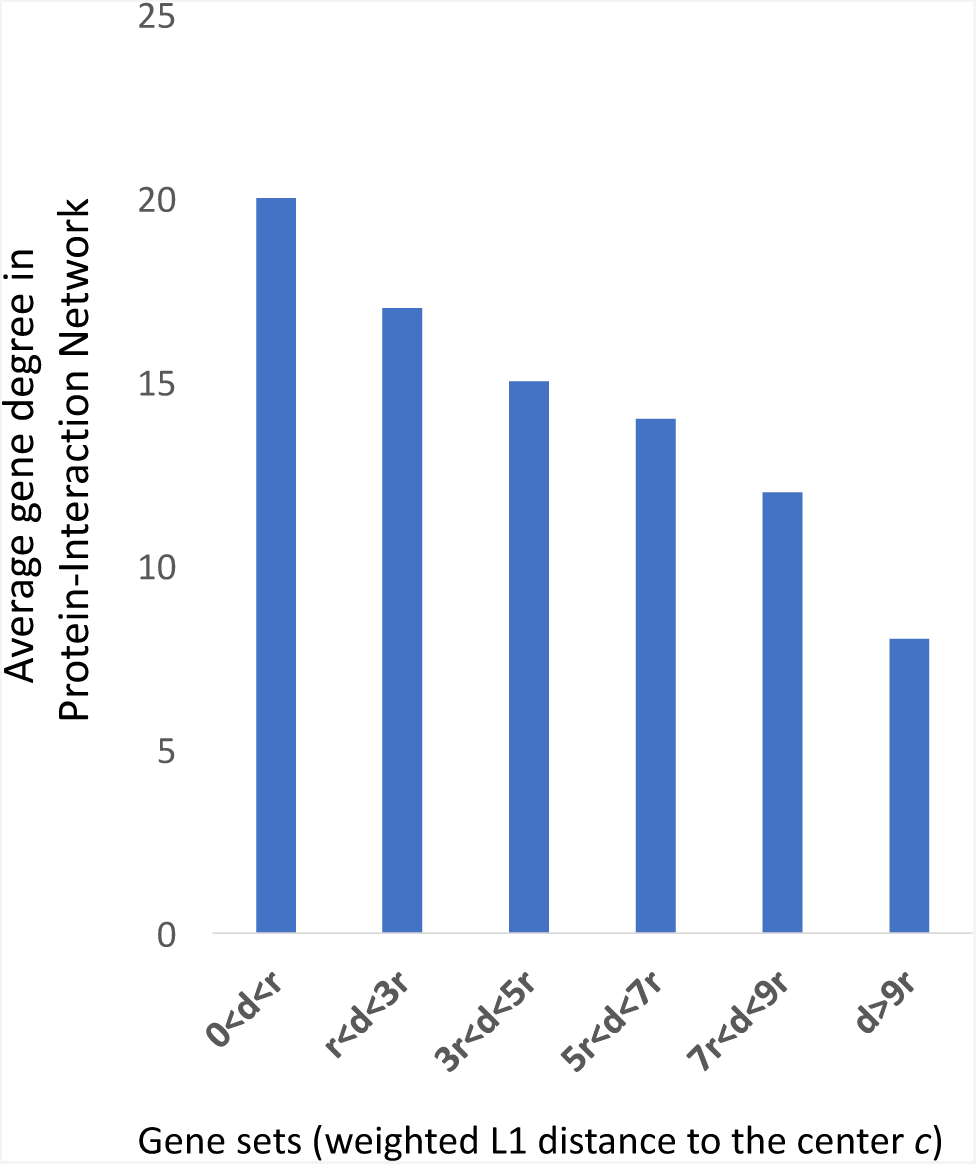
The average degree of genes is higher for set of genes which are closer to the center. The center and the weighted *ℓ*_1_ distance is learned by Odin.

### 1.5 Experiments details and commands

In the union of the ASD/ID datasets considered in this study (Table 1) there are a total of 684 affected ASD/ID cases/probands with LGD variants and 245 control and unaffected siblings with LGD variants. We compared the results of Odin against k-NN, SVM, and Glmnet (Lasso and Elastic-net) for predicting of ASD/ID with low false positive rate (< 1%). We used a leave-one-out approach to compare these methods. We used the scores/confidence/probability outputted by each method for each prediction to control for the number of unaffected samples predicted by mistake as case (denoted as false-positive rate). The exact commands used for each program is as follows:

#### SVM experiments

The command for training and testing used in for SVM is based on libSVM version 3.21 implementation [55]. Using the full dataset we first found the optimal parameters for “gamma” and “cost” and were set to 0.25 and 0.03125 respectively for the libSVM classifier. Then, for the LOO experiment we used the following commands in training dataset: `svm-train −b 1 −w0 5 −w1 1 −c 0.03125 −g 0.25 training-data` and in the case of test data we use the following command: `svm-predict −b 1 testing-data training-data.model output`.

#### Lasso and Elasticnet (Glmnet) experiments

The commands used for Glmnet (lasso and Elastic-net) [56]. In training dataset we use the following command: `fit=glmnet(training-data.features, training-data.class, alpha=a` (we ran with parameters *a* ∈ {0, 0.25, 0.5, 0.75, 1}, and in the case of test data we used `predict(fit, testing-data, s=0.042645)` (the value s was calculate as *lambda.min* as instructed in https://web.stanford.edu/hastie/glmnet/glmnetalpha.html).

#### K-NN experiments

We implemented the k-NN classier and tested and reported the results for *k* ranging from 1 to 20.

## Footnotes

† A preliminary version of this paper was accepted for presentation in RECOMB conference.

